# Targeting LINC00152 activates cAMP/Ca^2+^/ferroptosis axis and overcomes tamoxifen resistance in ER+ breast cancer

**DOI:** 10.1101/2023.11.05.565697

**Authors:** Ozge Saatci, Rashed Alam, Kim-Tuyen Huynh-Dam, Aynur Isik, Meral Uner, Nevin Belder, Pelin Gulizar Ersan, Metin Cetin, Unal Metin Tokat, Mustafa Emre Gedik, Hilal Bal, Ozlem Sener Sahin, Yasser Riazalhosseini, Denis Thieffry, Daniel Gautheret, Besim Ogretmen, Sercan Aksoy, Aysegul Uner, Aytekin Akyol, Ozgur Sahin

## Abstract

Tamoxifen has been the mainstay therapy to treat early, locally advanced, and metastatic estrogen receptor-positive (ER+) breast cancer, constituting around 75% of all cases. However, emergence of resistance is common, necessitating the identification of novel therapeutic targets. Here, we demonstrated that long-noncoding RNA LINC00152 confers tamoxifen resistance via blocking tamoxifen-induced ferroptosis, an iron-mediated cell death. Mechanistically, inhibiting LINC00152 reduces the mRNA stability of phosphodiesterase 4D (*PDE4D*), leading to activation of cAMP/PKA/CREB axis and increased expression of TRPC1 Ca^2+^ channel. This causes cytosolic Ca^2+^ overload and generation of reactive oxygen species (ROS) that is, on one hand, accompanied by downregulation of FTH1, a member of the iron sequestration unit, thus increasing intracellular Fe^2+^ levels; and on the other hand, inhibition of the peroxidase activity upon reduced GPX4 and xCT levels. These ultimately induce lipid peroxidation and ferroptotic cell death in combination with tamoxifen. Overexpressing PDE4D rescues LINC00152 inhibition-mediated tamoxifen sensitization by de-activating the cAMP/Ca^2+^/ferroptosis axis. Importantly, high LINC00152 expression is significantly correlated with high PDE4D/low ferroptosis and worse survival in multiple cohorts of tamoxifen- or tamoxifen-containing endocrine therapy-treated ER+ breast cancer patients. Overall, we identified LINC00152 inhibition as a novel mechanism of ferroptosis induction and tamoxifen sensitization, thereby revealing LINC00152 and its effectors as actionable therapeutic targets to improve clinical outcome in refractory ER+ breast cancer.

## Introduction

Estrogen receptor-positive (ER+) breast cancer accounts for almost 75% of all breast cancers. Endocrine therapies, modulating ER level and/or activity are the standard of care in ER+ breast cancer. The selective ER modulator (SERM), tamoxifen has been the backbone of endocrine therapy for over 40 years, especially for premenopausal women ^1,2^. Tamoxifen is recommended to treat early, locally advanced and metastatic ER-positive BCs^3^ and significantly improves overall survival^3,4^. Despite its initial clinical success, patients inevitably develop resistance and experience poor clinical outcome^5^. The molecular mechanisms of tamoxifen resistance are partially understood and involves ER downregulation, *ESR1* mutations, modulation of cell cycle regulators, activation of receptor tyrosine kinases and downstream effectors, and modulation of non-coding RNAs^6–10^. The clinical management of tamoxifen resistance involves the use of other endocrine therapies or CDK4/6 inhibitors^11^. However, resistance to the second-line therapies is also inevitable^12^. Therefore, there is still an unmet clinical need for identification of key processes and novel targets to overcome tamoxifen resistance and improve patient outcome.

Ferroptosis is a relatively newly discovered iron-dependent oxidative form of cell death that is characterized by lipid peroxidation, cell volume shrinkage and increased mitochondrial membrane density^13^. It is under tight control by the interplay between pro-oxidant and antioxidant systems^14^. Major indicators of ferroptosis are increase in lipid peroxidation and decreased expression or activity of the glutathione peroxidase 4 (GPX4), a major component of the antioxidant system^15^. Certain anti-cancer therapies have been shown to activate ferroptosis, such as sorafenib^16^, lapatinib^17^ and radiation therapy^18^, and vulnerability to ferroptotic cell death has been demonstrated to be a feature of certain therapy-resistant cancer cells^19^. Along these lines, ferroptosis induction is emerging as an effective strategy to eradicate therapy resistant cancer cells. However, the potential roles of ferroptosis in tamoxifen-mediated cell death or activating ferroptosis in overcoming resistance to endocrine therapies used in ER+ breast cancer, including tamoxifen, are mostly unknown.

Long non-coding RNAs (lncRNAs) are non-coding RNAs with more than 200 nucleotides in length that carry out diverse functions, including transcriptional regulation, regulation of proteins or RNA molecules by direct binding and stabilization, and sponging miRNAs^20^. Depending on the presence of specific features, they can be tissue- and/or disease-specific and may be detected in body fluids, thus making them highly attractive therapeutic targets and potential biomarkers^21^. LncRNAs may also drive resistance to anti-cancer therapies, including endocrine therapy via protecting ER against downregulation^22^, enhancing ER signaling^23^ or activating downstream PI3K/AKT or mTOR signaling^24^. LINC00152 is an oncogenic lncRNA, promoting survival, proliferation, EMT and invasiveness in cancer cells^25–27^. It has been associated with tumor aggressiveness^27,28^, and demonstrated to be a biomarker in various cancers, such as breast cancer, colorectal cancer and glioma^29–31^. Despite being a driver in several key oncogenic processes, the potential functions of LINC00152 in regulating distinct forms of cell death, such as ferroptosis to drive resistance to endocrine therapy, including tamoxifen in ER+ breast cancer remain to be elucidated.

Here we showed that LINC00152 is a driver of tamoxifen resistance, and its inhibition causes tamoxifen sensitization via reducing the mRNA stability of its novel interactor *PDE4D*, a cAMP-degrading phosphodiesterase. This leads to activation of cAMP/PKA/CREB axis which increases the expression of TRPC1 Ca^2+^ channel, causing cytosolic Ca^+2^ overload, generation of reactive oxygen species (ROS) and increase in lipid peroxidation, leading to ferroptosis in combination with tamoxifen. Notably, overexpressing PDE4D rescues LINC00152-mediated tamoxifen resistance by blocking ferroptosis.

## Results

### LINC00152 is upregulated in tamoxifen resistance and high LINC00152 expression correlates with worse prognosis

To identify differentially expressed lncRNAs involved in tamoxifen resistance, we performed RNA-sequencing (RNA-seq) using our previously generated acquired tamoxifen resistant (TamR) MCF-7 cells^32^ and identified 330 differentially expressed lncRNAs (q value = 0.05) between tamoxifen sensitive and resistant isogenic cells (**Supplementary Data 1**). LINC00152 was among the most significantly upregulated, highly abundant, validated and oncogenic lncRNAs in tamoxifen resistant cells compared to parental counterparts (**Fig. 1A, Supplementary Data 1**). We further validated its upregulation via qRT-PCR analysis in both MCF7 and T47D TamR cells compared to parental cells (**Fig. 1B**). To test the clinical relevance of LINC00152 upregulation in tamoxifen resistance, we first analyzed available gene expression profiling data of tamoxifen-treated ER+ breast cancer patients and demonstrated a significantly higher expression of LINC00152 in patients with no response to tamoxifen therapy compared to patients exhibiting tamoxifen response (**Fig. 1C**). A list of genes previously reported to be upregulated in tamoxifen resistance was enriched in ER+ breast cancer patients expressing higher LINC00152 (**Supplementary Fig. S1A**). To test the effects of LINC00152 levels on clinical outcome, we analyzed three independent datasets of tamoxifen-treated ER+ breast cancer patients and demonstrated that higher LINC00152 expression is significantly associated with worse disease-free and overall survival (**Fig. 1D-F**). We then performed in-situ hybridization (ISH) of LINC00152 in our own cohort of endocrine therapy-treated ER+ breast cancer patients and demonstrated that higher LINC00152 levels are associated with significantly worse disease-free survival (**Fig. 1G, H**). The LINC00152 ISH score was also higher in tumors with high Ki67 score and high grade (**Supplementary Fig. S1B, C**). Overall, we identified LINC00152 as a clinically relevant lncRNA upregulated in tamoxifen resistance.

**Figure 1.**
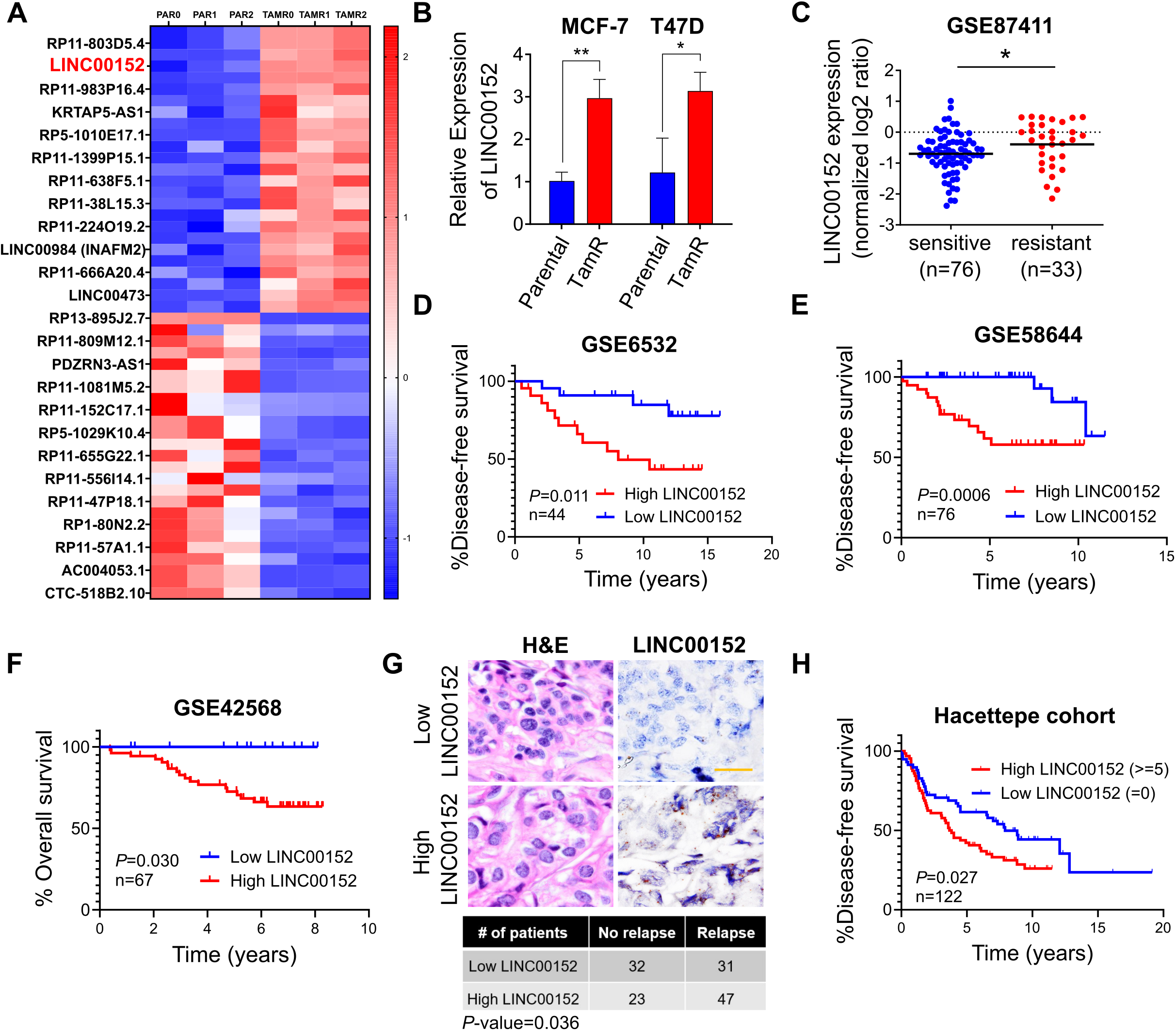
LINC00152 is upregulated in tamoxifen resistance and its higher expression correlates with poor prognosis. **A** Heatmap of the z-scores of the FPKM values of top 25 most abundant, upregulated, and top 25, least abundant, downregulated lncRNAs in MCF-7 tamoxifen resistant (TAMR) cells compared to parental (PAR) cells in three biological replicates (0, 1, 2). The names of 25 genes are depicted on the heatmap. **B** qRT-PCR analysis of LINC00152 in both MCF-7 and T47D parental and TamR cells. **C** LINC00152 levels in tamoxifen sensitive vs. resistant tumors at baseline from GSE87411. **D, E** The disease-free survival in tamoxifen-treated ER+ breast cancer patients with tumors expressing low vs. high LINC00152 expression from GSE6532 (D) and GSE58644 (E). **F** The overall survival in patients with tumors expressing low vs. high LINC00152 expression from GSE42568. **G** Representative *in situ* hybridization (ISH) staining of LINC00152 in low vs. high expressers among endocrine therapy-treated ER+ breast carcinoma specimens from the Hacettepe cohort. Scale bar = 20 µm. The numbers of Hacettepe patients having low vs high LINC00152 and with recurrence vs. no recurrence are provided at the bottom of the panel. **H** The disease-free survival in Hacettepe patients with tumors expressing low vs. high LINC00152 expression as quantified by ISH staining. Data are presented as mean values□±□standard deviation (SD). *P*-values for the bar graphs and boxplots were calculated with the unpaired, two-tailed Student’s t test. Significance for the Kaplan Meier survival graphs was calculated with Log-rank test. Chi-square test was used for G. * *P* < 0.05. Source data for this figure are provided as a Source Data file.

### Inhibiting LINC00152 overcomes tamoxifen resistance via inducing ferroptosis

To determine the roles of LINC00152 in tamoxifen resistance, we first determined the cellular localization of LINC00152. Cellular fractionation assay revealed that LINC00152 is primarily a cytoplasmic lncRNA (**Fig. 2A**). Here, lncRNAs DANCR^33^ and MALAT1^34^ are shown as positive controls, as there are known to be cytoplasmic and nuclear, respectively (**Fig. 2A**). To test the role of LINC00152 in tamoxifen resistance, we generated stable LINC00152 knockdown in MCF-7 TamR cells (MCF-7 TamR.shLINC00152) (**Fig. 2B**). LINC00152 knockdown led to a significant increase in cell viability inhibition in combination with tamoxifen in the MCF-7 TamR cells (**Fig. 2C**). These results were also recapitulated by using two different siRNAs against LINC00152 (**Supplementary Fig. S2A,** **Fig. 2D**). On the contrary, overexpressing LINC00152 led to significantly higher viability of the parental cells and reduced the viability inhibition induced by tamoxifen (**Supplementary Fig. S2B,** **Fig. 2E**).

**Figure 2.**
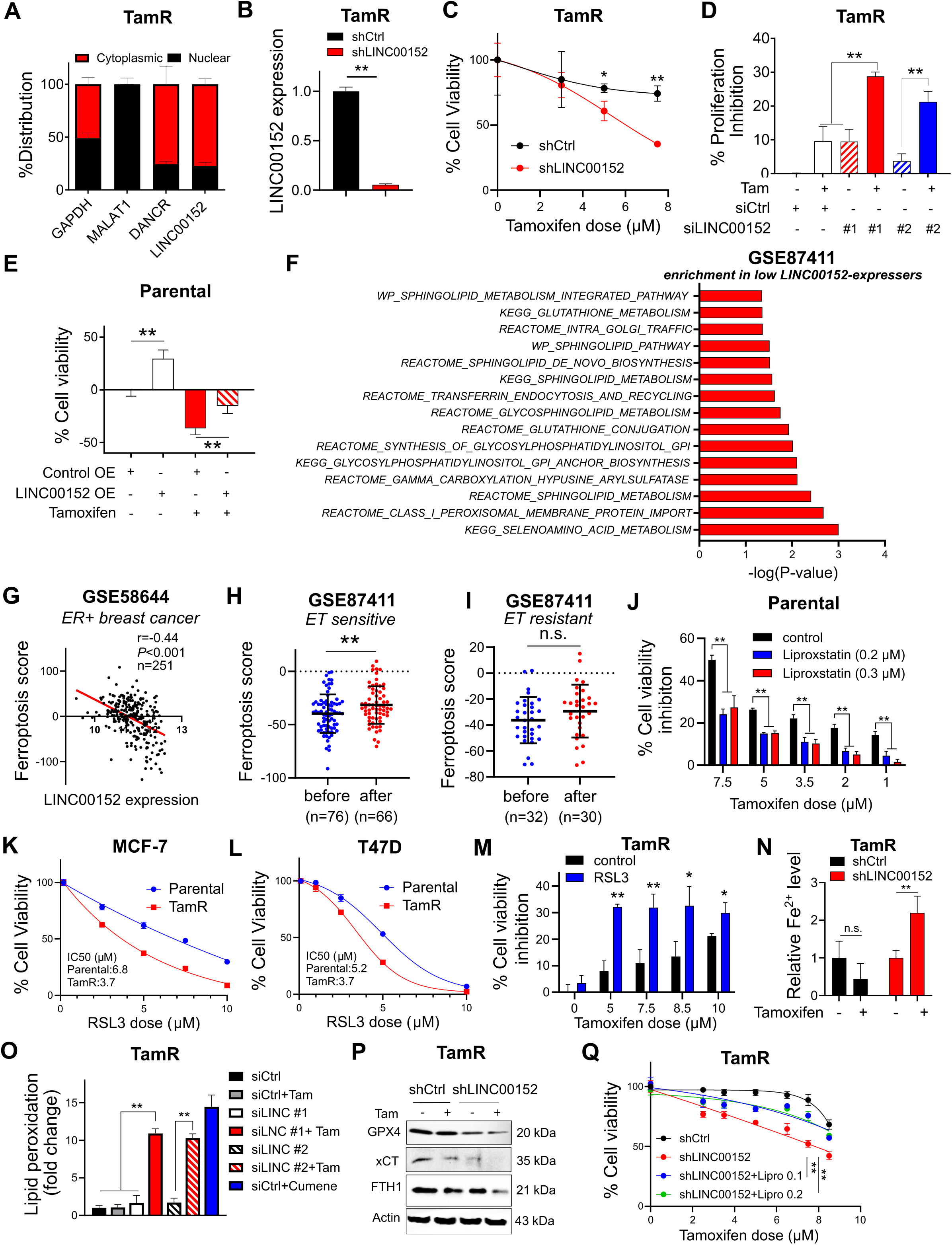
Inhibiting LINC00152 overcomes tamoxifen resistance by inducing ferroptosis. **A** The nuclear vs. cytoplasmic localization of LINC00152 in MCF-7 TamR cells. GAPDH, MALAT1 and DANCR were used as controls for nuclear and cytoplasmic RNAs. **B** LINC00152 knockdown efficacy in shLINC00152-expressing MCF-7 TamR cells. **C** Percent cell viability of MCF-7 TamR cells expressing shLINC00152 upon treatment with tamoxifen. **D** Percent viability inhibition in MCF-7 TamR cells transfected with siLINC00152 and treated with tamoxifen. **E** Viability assay in LINC00152-overexpressing T47D cells under tamoxifen treatment. **F** GSEA analysis showing the gene sets enriched in low LINC00152-expressing ER+ breast cancer patients treated with endocrine therapy from GSE87411. **G** Pearson correlation analysis of LINC00152 expression with ferroptosis score in ER+ patients from GSE58644. **H, I** Levels of ferroptosis score before and after endocrine therapy in sensitive (H) and resistant (I) ER+ patients from GSE87411. **J** Percentage cell viability inhibition in MCF-7 parental cells treated with tamoxifen with or without 0.2 or 0.3 µM of the ferroptosis inhibitor, liproxstatin-1. **K, L** Dose-response curves in MCF-7 (K) and T47D (L) parental vs. TamR cells treated with increasing doses of RSL3. **M** Percentage cell viability inhibition in MCF-7 TamR cells treated with tamoxifen alone or in combination with RSL3. **N** Relative intracellular Fe^2+^ levels in MCF-7 TamR shCtrl vs. shLINC00152 cells treated with 7.5 µM tamoxifen for 1.5 hrs. **O** Lipid peroxidation assay in T47D TamR cells transfected with siLINC00152 (siLINC) in combination with tamoxifen, showing fold change of lipid peroxidation. Cumene was used as a positive control. **P** Western blot analysis of the negative regulators of ferroptosis, xCT, GPX4 and FTH1 in MCF-7 TamR cells expressing shLINC00152 and treated with tamoxifen. Actin is used as a loading control in all Western blots unless stated otherwise. **Q** Percentage cell viability of shLINC00152-expressing MCF-7 TamR cells treated with tamoxifen with or without 0.1 or 0.2 µM of liproxstatin-1 (Lipro). Data are presented as mean values□±□standard deviation (SD). *P*-values for F were calculated using the GSEA software (Broad Institute) while others were calculated with the unpaired, two-tailed Student’s t test. * *P* < 0.05, ** *P* < 0.01, n.s. not significant. Source data for this figure are provided as a Source Data file. Source data for this figure are provided as a Source Data file.

Next, to identify the molecular mechanisms of LINC00152-mediated tamoxifen resistance, we performed RNA-seq upon siRNA-mediated knockdown of LINC00152 in the MCF-7 TamR cells. We generated a LINC00152 knockdown (KD) score comprising of genes down or upregulated upon LINC00152 knockdown (0.67 > FC > 1.5, *P* < 0.1). Accordingly, patients who express high LINC00152 KD score are expected to express low levels of LINC00152. Among patients who express high LINC00152 KD score, the tamoxifen sensitivity-related genes that are downregulated in tamoxifen resistance were enriched, supporting the role of LINC00152 in tamoxifen resistance (**Supplementary Fig. S3A**). Then, we performed pathway enrichment analysis and observed a significant enrichment of genes involved in ferroptosis among LINC00152-modulated mRNAs (**Supplementary Fig. S3B**). To further validate the pathological relevance of these data, we performed gene set enrichment analysis (GSEA) among endocrine therapy-treated ER+ breast cancer patients separated based on LINC00152 expression. As a result, we observed significant enrichment of several pathways involved in glutathione metabolism, sphingolipid pathway and selenoamino acid metabolism which are all related to ferroptosis in patients with low LINC00152 expression (**Fig. 2F**). We observed a significant negative correlation between LINC00152 expression and a validated ferroptosis signature score, generated using FerrDb database^35^ among ER+ breast cancer patients from an independent dataset (**Fig. 2G**) which further validates the pathological relevance of LINC00152-ferroptosis connection. Furthermore, there was a significant increase in the ferroptosis score in sensitive tumors, but not in resistant ones, after treatment with endocrine therapy (**Fig. 2H, I**), suggesting that ferroptosis induction is associated with clinical endocrine therapy sensitivity.

To experimentally validate a potential association between tamoxifen sensitivity and LINC00152-mediated tamoxifen resistance with ferroptosis, we first tested whether tamoxifen-induced cell death involves ferroptosis induction. Inhibiting ferroptosis using liproxstatin-1^36^ reduced tamoxifen-mediated inhibition of cell viability in parental cells (**Fig. 2J**), suggesting that tamoxifen in part, induces ferroptosis as a mechanism of cell death. Next, we compared the sensitivity of parental vs. TamR cells to the GPX4-degrading ferroptosis inducing agent, RSL3. We observed lower IC50 of RSL3 in TamR cells compared to parental counterparts (**Fig. 2K, L**), suggesting that TamR cells exhibit increased sensitivity to ferroptosis induction upon GPX4 downregulation. To further test if ferroptosis induction could overcome tamoxifen resistance, we combined RSL3 with tamoxifen in TamR cells and observed higher inhibition of cell viability (**Fig. 2M**). Next, we asked if LINC00152 inhibition could activate ferroptosis as a mechanism of cell death in TamR cells. Indeed, we observed that knockdown of LINC00152 in combination with tamoxifen in T47D TamR cells resulted in significantly increased intracellular Fe^2+^ levels (**Fig. 2N**) that was followed by lipid peroxidation (**Fig. 2O**), and deactivation of the cystine/glutamate antiporter xCT and the glutathione peroxidase GPX4, the major antioxidant systems^15,37^ (**Fig. 2P**). Furthermore, in line with increased intracellular Fe2+, we observed a decrease in the levels of the ferritin heavy chain 1 (FTH1), the member of cytosolic iron storage protein complex, chelating iron molecules, thus playing an important antioxidant role^38,39^ (**Fig. 2P**). Notably, inhibiting ferroptosis using the ferroptosis inhibitor liproxstatin-1 partially rescued LINC00152 knockdown-mediated tamoxifen sensitization (**Fig. 2Q**), demonstrating the key role of ferroptosis induction in LINC00152 knockdown-mediated tamoxifen sensitization. Overall, these data suggest that ferroptosis could be a clinically relevant cell-death mechanism of tamoxifen and that inhibiting LINC00152 overcomes tamoxifen resistance in part via induction of ferroptosis.

### LINC00152 inhibition increases cytosolic Ca^2+^ by activating cAMP signaling

Increase in cytosolic Ca^2+^ has been shown to be a major regulator of ferroptosis^40^. In line with this, we found a significant positive correlation between the scores of validated Ca^2+^ signaling^41^ and ferroptosis^35^ signatures in post endocrine therapy-treated sensitive ER+ breast cancer patients (**Fig . 3A**). Ca^2+^ signaling score was also significantly upregulated after endocrine therapy only in sensitive, but not in resistant patients (**Fig. 3B, C**), similar to the ferroptosis score (**Fig. 2H, I**). To test whether there is a change in cytosolic Ca^2+^ levels upon LINC00152 knockdown in combination with tamoxifen, we utilized a GFP-labeled Ca^2+^ reporter and observed a significant increase in Ca^2+^ levels in siLINC00152-transfected MCF-7 TamR cells treated with tamoxifen (**Fig. 3D**). To identify the regulator of calcium signaling under the control of LINC00152, we analyzed the RNA-seq data of siLINC00152-transfected cells and found that TRPC1, a Ca^2+^ channel located on cell or endoplasmic reticulum (EnR) membrane and regulating the Ca^2+^ levels within the cytosol^42^, was found to be upregulated upon LINC00152 knockdown (**Supplementary Fig. S3C**). We validated the RNA-seq data by qRT-PCR and demonstrated that the expression of TRPC1 is increased at both mRNA and protein levels upon LINC00152 knockdown and further increased in combination with tamoxifen in TamR cells (**Fig. 3E, F**).

**Figure 3.**
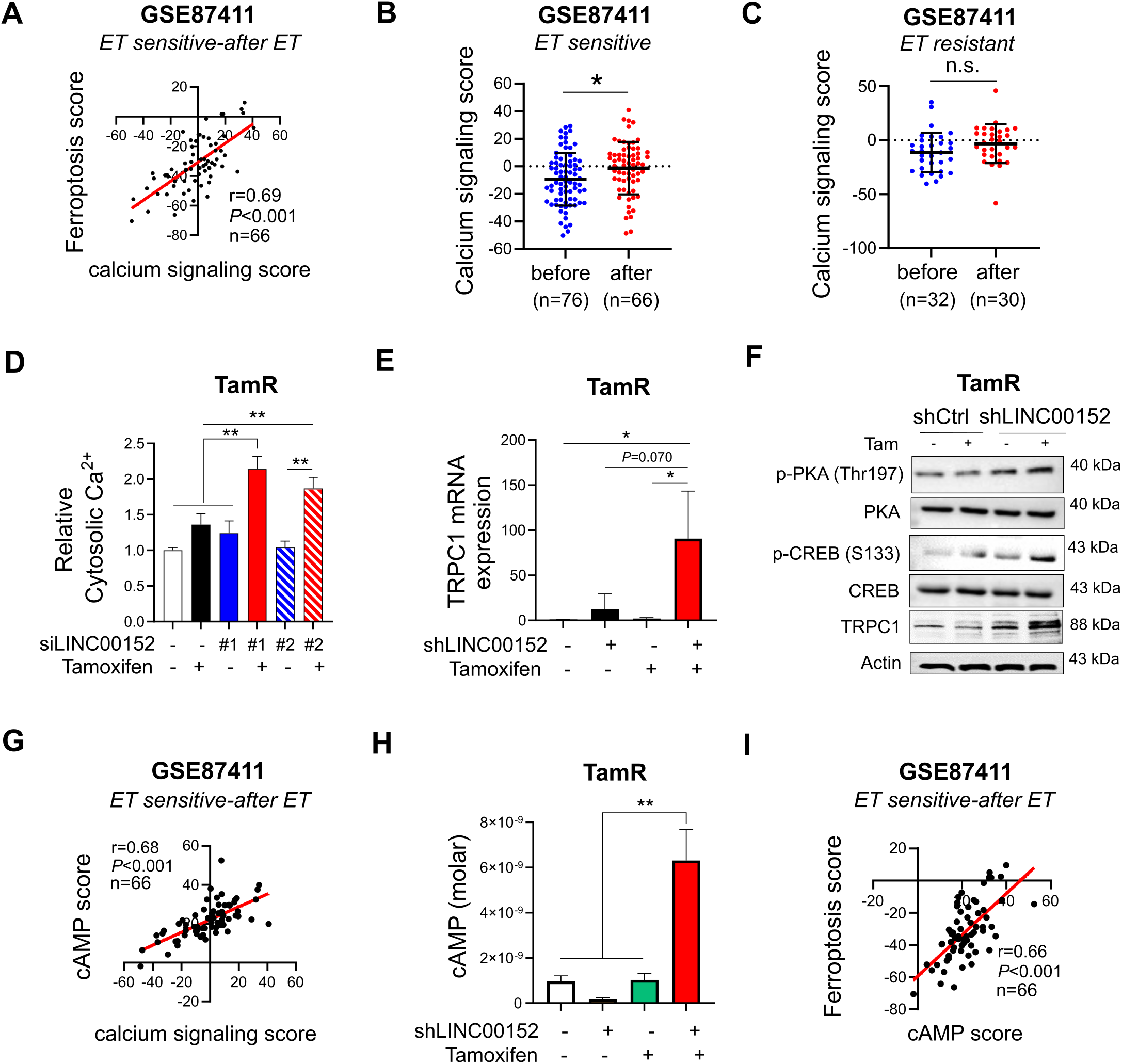
LINC00152 inhibition increases cytosolic Ca^2+^ by activating cAMP signaling. **A** The Pearson correlation analysis of calcium signaling with ferroptosis score in endocrine therapy sensitive ER+ patients after therapy from GE87411. **B, C** Levels of calcium signaling score before and after endocrine therapy in sensitive (B) and resistant (C) ER+ patients from GSE87411. **D** The levels of cytosolic Ca^2+^ in MCF-7 TamR cells transfected with two different siRNAs against LINC00152 and treated with tamoxifen. **E** qRT-PCR analysis of *TRPC1* mRNA in MCF-7 TamR shCtrl and shLINC00152 cells treated with tamoxifen. **F** Western Blot analysis of PDE4D, p-CREB (S133), p-PKA (Thr197) and TRPC1 in MCF-7 TamR shCtrl and shLINC00152 cells treated with tamoxifen. **G** The Pearson correlation analysis of calcium signaling score with cAMP score in endocrine therapy sensitive ER+ patients after therapy from GSE124647. **H** cAMP levels in MCF-7 TamR shCtrl or shLINC00152 cells treated with tamoxifen. **I** The Pearson correlation analysis of cAMP score and ferroptosis score in endocrine therapy sensitive ER+ patients after therapy from GSE87411. Data are presented as mean values□±□standard deviation (SD). *P*-values were calculated with the unpaired, two-tailed Student’s t test. * *P* < 0.05, ** *P* < 0.01, n.s. not significant. Source data for this figure are provided as a Source Data file.

To identify the molecular mechanisms of the LINC00152-regulated TRPC1 induction and cytoplasmic Ca^2+^ overload, we first examined a potential association between calcium and cAMP signaling since the increase in cAMP was shown to stimulate Ca^2+^ flux by activating the calcium channels^43^. We identified a significant positive correlation between calcium signaling score and cAMP signaling score^44^ in endocrine sensitive patients after treatment with endocrine therapy (**Fig. 3G**). We then experimentally measured cAMP levels and detected a significant increase in cAMP upon knocking down LINC00152 in combination with tamoxifen (**Fig. 3H**). The increase in cAMP levels was followed by phosphorylation of the downstream effectors, PKA and CREB, along with increased TRPC1 protein expression (**Fig. 3F**). Notably, levels of the cAMP score were significantly positively correlated with the ferroptosis score in post endocrine therapy-treated sensitive ER+ breast cancer patients (**Fig. 3I**), similar to calcium signaling score (**Fig. 3A**). Altogether, these data demonstrate that LINC00152 knockdown in combination with tamoxifen activates cAMP signaling and increases TRPC1 levels, thus triggering cytoplasmic Ca^2+^ overload that likely initiates ferroptosis in TamR cells.

### LINC00152 inhibits ferroptosis by regulating PDE4D and downstream cAMP/Ca^2+^ signaling in tamoxifen resistant cells

PDE4D is one of the cAMP-specific phosphodiesterases that degrades cAMP^45^. We found that *PDE4D* mRNA is downregulated in the RNA-seq data of siLINC00152-transfected MCF-7 TamR cells (**Supplementary Fig. S4A**). The reduction was further validated at both mRNA and protein levels by qRT-PCR and Western blotting, respectively, upon siRNA or shRNA-mediated LINC00152 knockdown in TamR cells (**Fig. 4A, B, Supplementary Fig. S4B, C**). Conversely, overexpressing LINC00152 in parental MCF-7 and T47D cells significantly increased the mRNA and protein expressions of PDE4D (**Supplementary Fig. S4D, E**). Notably, combining tamoxifen with the PDE4D inhibitor GebR-7b^46^ increased TRPC1 levels along with activation of PKA/CREB and reduction of GPX4 (**Fig. 4C**), phenocopying the effects of LINC00152 knockdown (**Fig 3F**). Next, we tested the clinical association of PDE4D with ferroptosis and demonstrated significantly lower ferroptosis score in patients expressing high levels of PDE4D (**Fig. 4D**).

**Figure 4.**
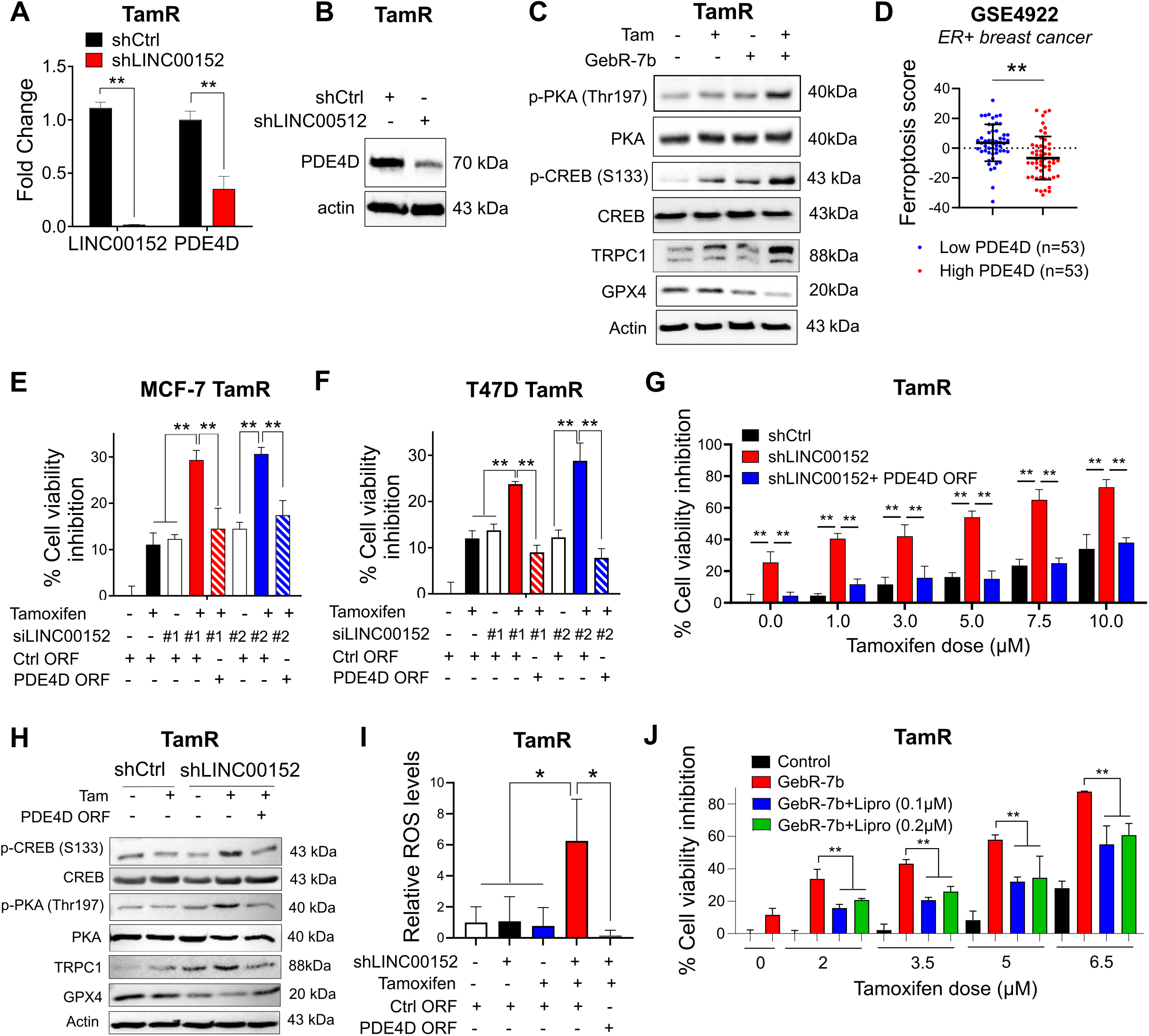
LINC00152 inhibits ferroptosis by regulating PDE4D and downstream cAMP/Ca^2+^ signaling in tamoxifen resistant cells. **A** qRT-PCR analysis of LINC00152 and PDE4D in the shLINC00152-expressing MCF-7 TamR cells. **B** Western blot analysis of PDE4D in the shLINC00152-expressing MCF-7 TamR cells. **C** Western blot analysis of p-CREB (S133), p-PKA (Thr197), TRPC1 and ferroptosis inhibitor GPX4 in T47D TamR cells treated with tamoxifen in combination with GebR-7b. **D** Levels of ferroptosis score in low vs. high PDE4D-expressing ER+ patients from GSE4922. **E, F** Percentage cell viability inhibition in MCF-7 and T47D TamR cells co-transfected with siLINC00152 and PDE4D ORF and treated with tamoxifen. **G** Percentage of viability inhibition in shLINC00152-expressing MCF-7 TamR cells stably overexpressing PDE4D and treated with tamoxifen. **H** Western blot analysis of p-CREB (S133), p-PKA (Thr197), TRPC1 and GPX4 in shLINC00152-expressing MCF-7 TamR cells stably overexpressing PDE4D and treated with tamoxifen. **I** Relative ROS levels in MCF-7 TamR cells expressing shLINC00152 and treated with tamoxifen. **J** Percentage viability inhibition in T47D TamR cells treated with tamoxifen and GebR-7b with or without 0.1 or 0.2 µM of liproxstatin-1 (Lipro). Data are presented as mean values□±□standard deviation (SD). *P*-values were calculated with the unpaired, two-tailed Student’s t test. * *P*< 0.05, ** *P* < 0.01. Source data for this figure are provided as a Source Data file.

To test the contribution of PDE4D downregulation upon LINC00152 knockdown to tamoxifen sensitization, we performed a rescue experiment where we overexpressed PDE4D ORF along with LINC00152 knockdown followed by tamoxifen treatment. As shown in **Fig. 4E-G**, PDE4D overexpression completely rescued cell viability in the presence of siRNA or shRNA-mediated LINC00152 knockdown in combination with tamoxifen. Notably, overexpressing PDE4D in the presence of shLINC00152 together with tamoxifen reduced TRPC1 expression along with de-activation of PKA and CREB and increased expression of GPX4 (**Fig. 4H**). Importantly, we detected a strong induction of reactive oxygen species (ROS), a crucial initiator of ferroptosis^39^, upon LINC00152 knockdown in combination with tamoxifen, which was reversed upon PDE4D overexpression (**Fig. 4I**). Along these lines, inhibiting ferroptosis using liproxstatin-1 partially rescued cell viability under treatment with tamoxifen and GebR-7b (**Fig. 4J**). Overall, these data suggest that LINC00152 blocks ferroptosis and mediates tamoxifen resistance via regulating PDE4D/cAMP/Ca^2+^ signaling axis.

### LINC00152 stabilizes *PDE4D* mRNA by interacting with its 3’UTR

lncRNAs can modulate gene expression in multiple ways, one of which is to regulate the stability of the target mRNA^47^. We observed a positive correlation between LINC00152 and *PDE4D* mRNA in ER+ breast cancer patients, suggesting mRNA level regulation of PDE4D by LINC00152 (**Fig. 5A**). PDE4D also showed a positive correlation with LINC00152 at the protein level in our own cohort of ER+ breast cancer patients (**Fig. 5B, C**). To test whether *PDE4D* mRNA stability is affected by LINC00152 modulation, we performed mRNA stability assay under actinomycin treatment in MCF-7 and T47D TamR cells transfected with siLINC00152. As shown in **Fig. 5D, E**, LINC00152 knockdown reduced the half-life of *PDE4D* mRNA in both MCF-7 and T47D TamR cells. To further confirm the specific interaction between LINC00152 and *PDE4D* mRNA, we performed RNA pull down and demonstrated the direct binding between LINC00152 and *PDE4D* mRNA (**Fig. 5F**). The 3’UTR region of mRNAs is usually responsible for regulating the stability of mRNAs^48^. Therefore, we reasoned that LINC00152 is potentially interacting with the 3’UTR region of the *PDE4D* mRNA to modulate its stability. To test this hypothesis, we first examined the predicted binding sites between *PDE4D* 3’UTR and LINC00152 using the IntaRNA tool^49^ and identified three potential interaction sites on the *PDE4D* 3’UTR (**Fig. 5G, Supplementary Fig. S5A, B**). We then cloned the 3’UTR of *PDE4D* in 3 different pieces, each containing one predicted binding site for LINC00152 (UTR#1, 2, 3) upstream of the luciferase gene. Both MCF-7 and T47D cell lines were co-transfected with *PDE4D* 3’UTR vectors together with LINC00152 overexpression vector. This resulted in an increase in luciferase signal from UTR#1 (**Fig. 5H, I**), suggesting that LINC00152 potentially binds to 3’UTR region#1 (binding energy: -18.56 kcal/mol, **Fig. 5G**) and increases *PDE4D* mRNA stability. Overall, these data demonstrate that LINC00152 interacts with *PDE4D* 3’ UTR in TamR cells, increasing its mRNA stability, and thereby mediates tamoxifen resistance.

**Figure 5.**
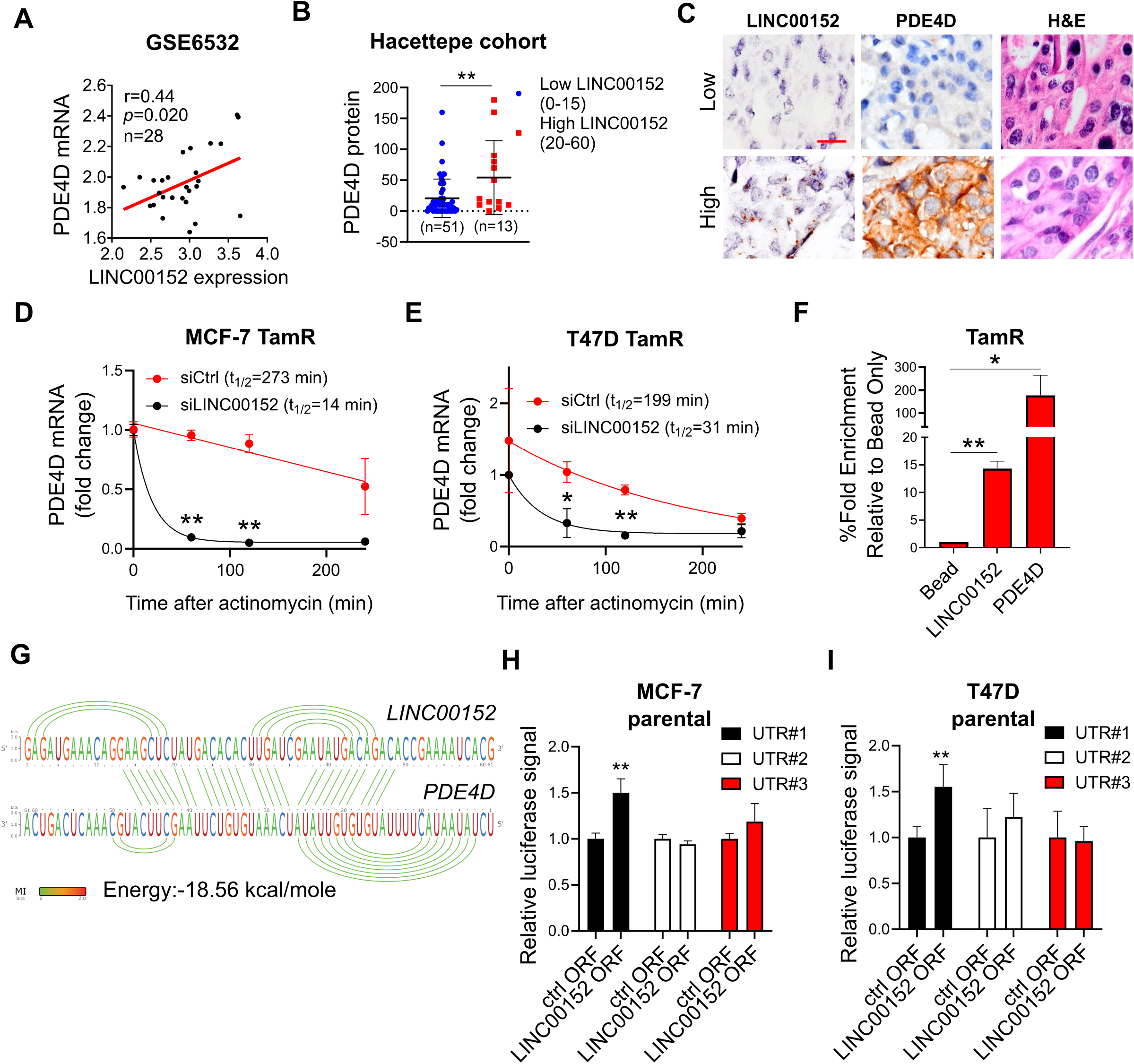
LINC00152 stabilizes *PDE4D* mRNA by interacting with its 3’UTR. **A** The Pearson correlation analysis of LINC00152 and *PDE4D* mRNA in patients from GSE6532. **B** PDE4D protein expression determined by IHC in low vs. high LINC00152-expressing patients from the Hacettepe cohort. **C** Representative images of LINC00152 ISH, PDE4D IHC and H&E staining in patients from the Hacettepe cohort. Scale bar = 20 µm. **D, E** qRT-PCR analysis of PDE4D in MCF-7 TamR (D) and T47D TamR (E) cells transfected with siLINC00152 for 48 hours followed by treatment with 5 µg/ml of Actinomycin D. **F** Percentage fold enrichment of PDE4D in LINC00152 pull down in MCF-7 TamR cells. **G** Interaction between LINC00152 and *PDE4D* 3’UTR predicted by the IntaRNA tool. The visualization of the interaction was obtained using RILogo. **H, I** Luciferase assay in parental MCF7 (H) and T47D (I) cells co-transfected with LINC00152 overexpression vector and *PDE4D* 3’UTR, demonstrating binding to *PDE4D* 3’UTR. Data are presented as mean values□±□standard deviation (SD). *P*-values were calculated with the unpaired, two-tailed Student’s t test. * *P* < 0.05, ** *P* < 0.01. Source data for this figure are provided as a Source Data file.

### LINC00152 inhibition overcomes tamoxifen resistance in vivo

To test the effects of LINC00152 knockdown on in vivo tamoxifen sensitization, we first analyzed the expression of LINC00152 in MCF-7 TamR vs. parental tumors grown in mice and observed a significant upregulation of LINC00152 also within in vivo-grown MCF-7 TamR xenografts (**Fig. 6A**) similar to cultured cells (**Fig. 1B**). We then generated xenografts of MCF-7 TamR.shCtrl and shLINC00152 cells and treated the mice with vehicle (corn oil) or tamoxifen (3 mg/kg, daily) for a month. As shown in **Fig. 6B-D**, there was a significant reduction of tumor growth and tumor weight upon LINC00152 knockdown in combination with tamoxifen. qRT-PCR analysis of LINC00152 and immunohistochemistry (IHC) staining of PDE4D in tumors collected in the end of the experiment demonstrated that LINC00152 knockdown was sustained in vivo (**Fig. 6E**) and PDE4D expression was strongly downregulated (**Fig. 6F**), validating our in vitro findings. Overall, we showed that LINC00152 inhibition overcame tamoxifen resistance in vivo, at least in part, via downregulating PDE4D expression.

**Figure 6.**
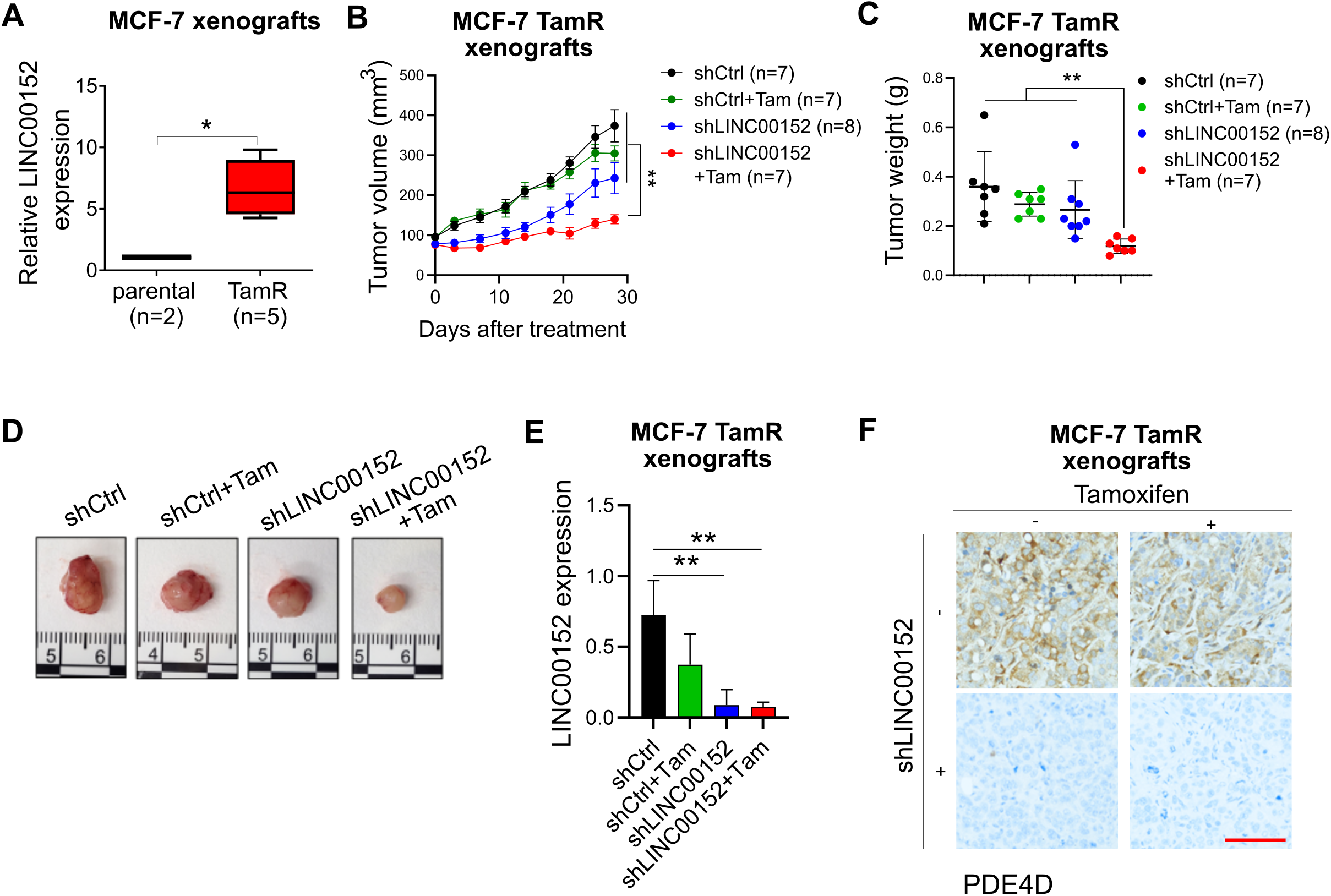
LINC00152 inhibition overcomes tamoxifen resistance in vivo. **A** *PDE4D* mRNA expression in MCF-7 parental (PAR) vs. TamR xenograft tumors. **B** Tumor volumes of MCF-7 TamR xenografts expressing shLINC00152 and treated with tamoxifen (3 mg/kg, daily). **C, D** Tumor weights (C) and tumor images (D) from mice in B. **E** qRT-PCR analysis of LINC00152 in tumors from B. **F** IHC staining of PDE4D in tumors from B. Scale bar = 100 µm. Data for the bar graphs and box plots are represented as mean values□±SD, while data for the tumor volume graph are represented as mean values□± standard error of the mean (SEM). *P*-values for the bar graphs and box plots were calculated with the unpaired, two-tailed Student’s t test. Significance for the tumor volume graph was calculated with two-way ANOVA. Source data for this figure are provided as a Source Data file.

## Discussion

Tamoxifen is the first ER modulator approved by the FDA in 1970s and it has been the backbone of adjuvant hormone therapy^2^. However, resistance develops in approximately 20-30% of high risk, advanced ER-positive breast cancer patients, decreasing survival^50^. Therefore, there is a dire need to identify novel therapeutic strategies to improve the clinical outcome for patients with resistance to tamoxifen. In this study, we demonstrated that ferroptosis is involved in tamoxifen-mediated cell death, and tamoxifen resistance is characterized by higher vulnerability to ferroptosis. We identified LINC00152 as a driver of tamoxifen resistance, which enhances the mRNA stability of its novel target *PDE4D*, thus de-activating cAMP/PKA/CREB axis and blocking ferroptosis (**Fig. 7A**). On the other hand, LINC00152 inhibition destabilizes PDE4D and activates cAMP/PKA/CREB, similar to targeting PDE4D with GebR-7b, and increases TRPC1, cytosolic Ca^2+^, ROS and Fe^2+^ levels, accompanied by reduced levels of the antioxidant systems FTH1, GPX4, and xCT, leading to lipid peroxidation, ferroptotic cell death and tamoxifen sensitization (**Fig. 7B**). Notably, we showed that high LINC00152 expression is significantly correlated with high PDE4D/low ferroptosis and worse survival in multiple cohorts of tamoxifen- or tamoxifen-containing endocrine therapy-treated ER+ breast cancer patients.

**Figure 7.**
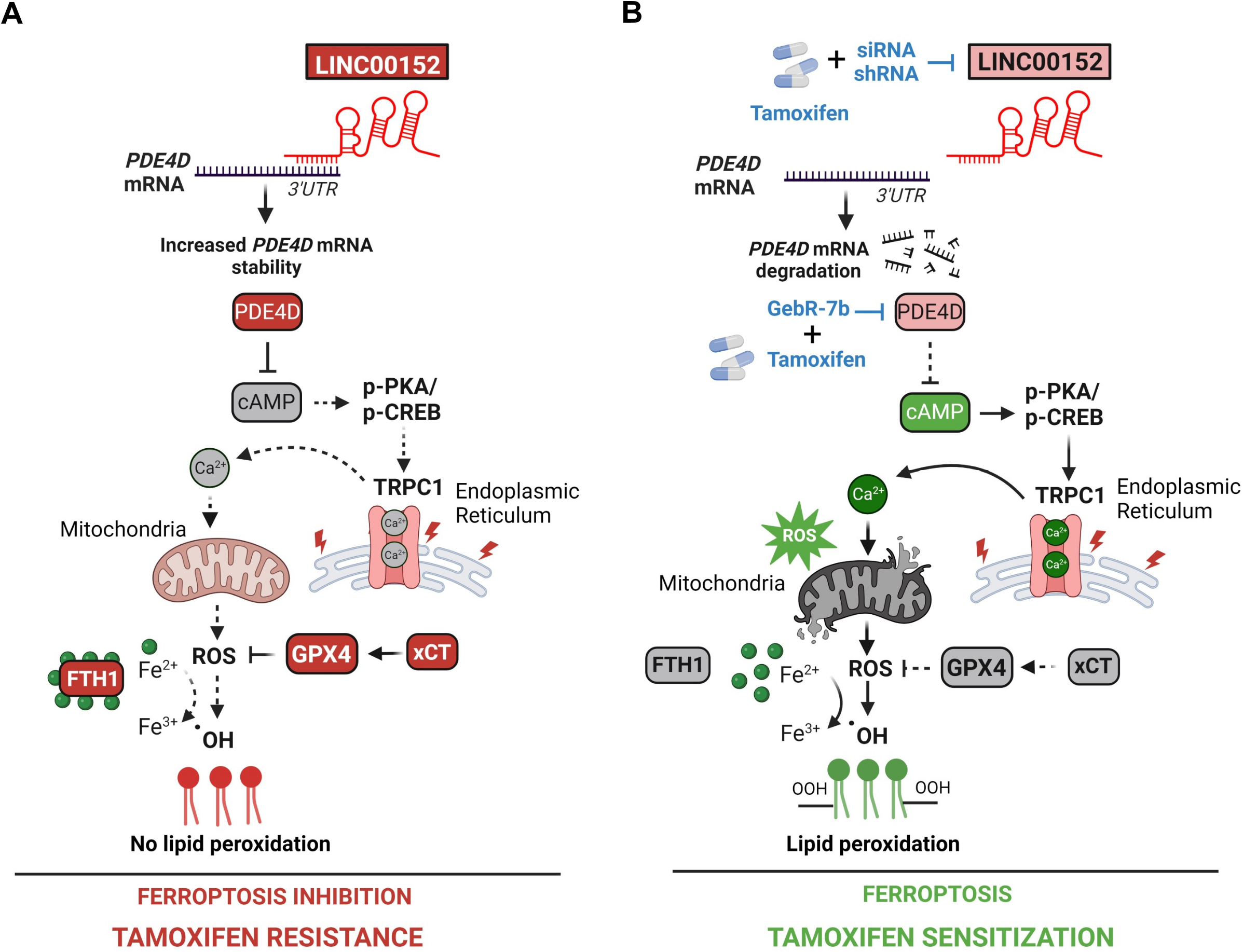
Schematic summary of LINC00152/PDE4D-driven tamoxifen resistance. **A** LINC00152 is overexpressed in tamoxifen resistance, binds to *PDE4D* 3’UTR, and increases its mRNA stability. This results in reduced cAMP levels, and de-activation of PKA/CREB, leading to decreased expression of TRPC1, reduction of cytosolic Ca^2+^ and blockage of reactive oxygen species (ROS) generation. Furthermore, the negative regulators of ferroptosis, FTH1, GPX4 and xCT are upregulated, and contribute to reduction of intracellular Fe^2+^, and inhibition of lipid peroxidation that ultimately block ferroptosis-mediated cell death and mediate tamoxifen resistance. **B** Targeting LINC00152 in combination with tamoxifen leads to reduced mRNA stability of PDE4D due to loss of binding to its 3’UTR, thereby activating cAMP/PKA/CREB, similar to targeting PDE4D with GebR-7b in combination with tamoxifen. This further increases TRPC1, elevates cytosolic Ca^2+^, generates reactive oxygen species (ROS) and increases Fe^2+^, accompanied by reduced levels of FTH1, GPX4 and xCT, leading to lipid peroxidation, ferroptotic cell death and tamoxifen sensitization.

Activation of ferroptosis is initiated by an imbalance between pro-oxidant and antioxidant systems^14^ and is emerging as an attractive strategy to overcome resistance to certain drugs, such as cisplatin ^51,52^. The ROS-mediated lipid peroxidation is one of the key steps driving ferroptosis upon oxidative degradation of lipids and the subsequent membrane damage^39^. On the other hand, xCT and GPX4 inhibit ferroptosis by promoting the synthesis of reduced glutathione (GSH), the primary antioxidant molecule^37^, and reducing phospholipid hydroperoxide production^15^, respectively. The current ferroptosis-inducing strategies used to eliminate therapy-resistant cancer cells usually target GPX4 by downregulating its expression^53^ or by inhibiting its activity via the reduction of its cofactor, GSH^54^, thus disturbing the balance between oxidant and antioxidant systems. Here, we showed that our novel combination of tamoxifen and LINC00152 knockdown induces ferroptosis by not only deactivating the antioxidant defense via reducing xCT and GPX4, but also by increasing Ca^2+^ overload through modulating the PDE4D/cAMP/PKA/CREB axis, thus causing ROS generation, accompanied by loss of the iron chelator, FTH1 that ultimately increases Fe^2+^ levels, and results in lipid peroxidation. Our data showing higher sensitivity to ferroptosis induction in TamR cells is also supported by the gain of mesenchymal traits in tamoxifen resistance, previously shown by us^55^ and others^56,57^, since epithelial to mesenchymal transition (EMT) has recently been shown to increase vulnerability to ferroptosis sensitivity^58^. Furthermore, we recently showed that endocrine therapy and CDK4/6 inhibitors commonly increase ROS levels in cancer cells and leads to cAMP-induced DNA damage, leading to PARP trapping and transcriptional blockage^59^. Future studies may investigate whether sensitivity to ferroptosis-mediated cell death is altered also in resistance to other endocrine therapies or CDK4/6 inhibitors, and whether LINC00152 targeting could also be used as an effective strategy to overcome resistance to these agents, or in other resistant models vulnerable to ferroptosis induction.

Elevated levels of second messengers, such as cAMP and Ca^2+^, are known to be involved in stress response, controlling cell survival and/or death^60,61^. Endoplasmic reticulum (EnR) is the center for managing cellular stress^62^ and is the main storage site for Ca^+2^ ^63^. A few anti-cancer agents are known to mediate cell death via activating EnR stress, such as bortezomib^64^ and cisplatin^65^. However, the mechanisms of stress response induction in endocrine resistance are mostly undiscovered. We previously reported a novel mechanism of tamoxifen sensitization via inducing EnR stress, leading to activation of p38/JNK signaling and apoptosis in tamoxifen resistant ER+ breast cancer cells^32^. Here, we characterized yet another novel type of stress response leading to tamoxifen sensitization, involving Ca^2+^ overload upon LINC00152/PDE4D inhibition. Recent evidence suggests that manipulating Ca^2+^ homeostasis stands as an attractive strategy to restore ferroptosis sensitivity. For instance, a novel tetraspanin, MS4A15 located on EnR membrane blocks ferroptosis via depleting luminal Ca^2+^ stores and reprogramming lipids^66^. We identified TRPC1, a Ca^2+^ channel that can be located on cell or EnR membrane^42^, as a potentially novel regulator of ferroptosis. We showed that TRPC1 is upregulated upon activation of cAMP/PKA/CREB, increases cytosolic Ca^2+^ and ROS levels. Importantly, EnR is known to be in crosstalk with mitochondria^67^, and an excessive flux of Ca^2+^ from EnR to mitochondria can lead to increased respiratory chain activity and production of excessive ROS^68^, supporting our findings. Although TRPC1 has not been reported as a CREB-induced gene, knockdown of CREB was shown to downregulate one of the G-protein coupled dopamine receptors responsible for Ca^2+^ influx, thereby reducing calcium signaling^68^. This raises the possibility that TRPC1 could also be transcribed in a CREB-dependent manner upon downregulation of LINC00152/PDE4D. Furthermore, the complete rescue of cell viability upon overexpressing the PDE4D, in the presence of LINC00152 knockdown and tamoxifen, by reduction of TRPC1 and blockage of ROS, suggest that PDE4D is a novel and highly crucial modulator of ferroptotic cell death, and that targeting PDE4D can further be tested to enhance sensitivity to ferroptosis-mediated cell death.

We demonstrated the clinical relevance of the LINC00152/PDE4D/cAMP/Ca^2+^/ferroptosis axis and showed that higher LINC00152 levels are associated with worse survival in multiple cohorts of endocrine therapy-treated ER+ breast cancer patients. These results suggest that LINC00152 could be a potential biomarker for prognosis. However, future studies are warranted to test its sensitivity and specificity with larger cohorts of patients. In addition to being a potential biomarker, LINC00152 could also be targeted in clinics. LncRNAs are highly attractive targets for cancer therapy given their tissue specific expression, thus enabling specific targeting of tumors^69^. Importantly, in the last decade, RNA-based therapies, such as antisense oligonucleotides (ASOs) or siRNAs have undergone substantial improvements and are now being tested in clinical trials for various diseases, including cancer^70^. Overall, our data not only identify a novel mechanism of tamoxifen sensitization involving the regulation of ferroptosis by LINC00152, but also demonstrate the potential of LINC00152 as a clinically relevant resistance mediator, thus encouraging the clinical testing of LINC00152 targeting to overcome tamoxifen resistance.

## Materials and Methods

### Cell lines, drugs, and culture conditions

Human breast cancer cell lines, MCF-7 and T47D were purchased from ATCC and were cultured in phenol red–free DMEM (Gibco) with 10% FBS, 0.1% insulin, 50 U/mL penicillin/streptomycin, 1% nonessential amino acids (Gibco). Tamoxifen resistant (TamR) MCF-7 and T47D cells were generated previously^32^. Tamoxifen and GebR-7b were purchased from Sigma. Liproxstatin-1 and RSL3 were purchased from MedChem Express. Cells were routinely tested for mycoplasma contamination using MycoAlert detection kit (Lonza) and were authenticated by STR sequencing.

### ER+ human tumor samples

To analyze the effects of LINC00152 expression on the survival of endocrine therapy-treated female ER+ breast cancer patients, and its correlation with PDE4D protein, we performed in situ hybridization (ISH) of LINC00152 and immunohistochemistry (IHC) staining of PDE4D in primary tumor samples from 147 ER+ breast cancer patients that were diagnosed between 2001 and 2018 at Hacettepe University School of Medicine, Ankara, Turkey and treated with endocrine therapy with or without radiotherapy and chemotherapy. The study was approved by the Non-Interventional Clinical Research Ethics Committee of Hacettepe University (approval no: 2020/02-40).

### Whole-transcriptome sequencing (RNA-seq) and data analysis

RNA-sequencing of MCF-7 parental vs. TamR cells was performed in triplicates using the Illumina HiSeq 2000 platform at McGill University and Genome Quebec Innovation Centre as described previously^32^. The 330 differentially expressed lncRNAs between parental vs. TamR cells (q value = 0.05) were retrieved **(Supplementary Data 1**). RNA-sequencing of MCF-7 TamR cells transfected with siControl, siLINC00152#1 or siLINC00152#2 was performed for each condition in duplicates at University of South Carolina Functional Genomics Core. The genes differentially expressed upon LINC00152 knockdown (siControl vs. siLINC00152#1 and/or siControl vs. siLINC00152#2) with the cut-off of 0.67 > FC > 1.5 and *P* < 0.1 were determined. The details are provided in Supplementary Methods.

### Lipid peroxidation assay

For measuring the levels of lipid peroxidation as an indicator of ferroptosis, Image-iT™ Lipid Peroxidation Kit (ThermoFisher) was utilized following manufacturer’s protocol. Cells that were stained with the BODIPY™ 581/591 C11 dye provided in the kit for 30 minutes were analyzed by flow cytometry. The shift from red to green signal that occurs upon oxidation by lipid hydroperoxides were quantified by taking the ratio of the mean fluorescence emission at ∼ 590 nm to ∼510 nm and plotted as the relative ferroptosis levels.

### Cytosolic Ca^2+^ measurement

To detect cytosolic Ca^2+^ levels, the MCF-7 TamR cells were co-transfected with two different siLINC00152 and the GFP-based Ca^2+^ indicator, GCaMP3 (Addgene#64853) in 96-well format. One day later, cells were treated with 10 µM tamoxifen for 10 min and the fluorescence signal was measured with SpectraMax microplate reader.

### RNA pulldown

T47D TamR cells were lysed in freshly prepared lysis buffer with the addition of RNase inhibitor and Protease inhibitors (20 mM Tris-HCl, pH 7.4, 200 mM KCl, 150 mM NACl, 1% Triton-X-100, 0.1% SDS, 1 mM DTT, protease inhibitor cocktail and 200 U/mL of Riboblock). 1 mg of total lysates were mixed with 100 pmol of biotinylated probe against LINC00152 (**Supplementary Table S1**) and incubated for 1 hour at room temperature. Then, the lysates were incubated with blocked streptavidin beads and incubated overnight at 4°C. Next day, beads were incubated with Proteinase K buffer (10 mM Tris-HCl pH 7.0, 100 mM NaCl, 1 mM EDTA, 0.5% SDS) and 5 μL of proteinase-K (20 mg/mL) at 56°C for 1 hour and boiled at 95°C for 5 minutes. RNA was extracted via RNA purification kit (Zymo, R1013), converted to cDNA and analyzed by RT-qPCR using the primers listed in **Supplementary Table S2**. %Input method was used to calculate enrichment relative to input samples and normalized to only bead control.

### Intracellular cAMP measurements

To measure intracellular cAMP levels, MCF-7 TamR.shLINC cells that were treated with tamoxifen for 30 min were washed once with PBS and detached using Tryple. 3,000 cells/well were collected in eppendorfs and lysed with ice-cold 2.5 mol/L perchloric acid followed by neutralization with 2 M KOH. Cells were incubated on ice for 30 minutes and then centrifuged at 10,000 rpm for 10 min to collect the clear lysate. Twenty microliters of the clear lysate were transferred per well of a white plate, and cAMP measurement was done using the LANCE Ultra cAMP detection kit (PerkinElmer) according to the manufacturer’s protocol.

### Intracellular ROS measurement

The intracellular ROS was measured using the DCFDA / H2DCFDA - Cellular ROS Assay Kit from Abcam according to the manufacturer’s protocol. Briefly, 8,000 MCF-7 TamR.shCtrl and shLINC00152 cells were seeded on clear bottom, black wall plates and next day transfected with PDE4D ORF. One day later, cells were treated with tamoxifen for 9 hours flowed by staining with 20 µM DCFDA for 1 hour. Then, the fluorescence was detected by a Multimode SpectraMax plate reader at 37 ^0^C with excitation/emission at 485 nm/535 nm. The background GFP signal of the cells and the media were subtracted, and ROS levels were represented as relative change to shCtrl cells without any treatment.

### Intracellular Fe^2+^ measurement

The intracellular Fe^2+^ levels were measured using the FerroOrange Assay Kit from Dojindo Molecular Technologies (JAPAN) (F374) according to the manufacturer’s protocol. Briefly, MCF-7 TamR shCtrl vs. shLINC00152 cells were seeded on clear bottom, black wall plates and next day treated with 7.5 µM tamoxifen. The FerroOrange dye was added to media at a 1:1000 dilution immediately after treatment. After 1.5 hours, the fluorescence signal was quantified at Ex/Em = 561nm/600nm. The signal from only media was subtracted from all measurements and the relative Fe^2+^ levels were calculated by taking the ratio of tamoxifen-treated shCtrl or shLINC00152 cells to their corresponding untreated counterparts.

### In vivo mice experiments

All the in vivo studies were carried out in accordance with the Institutional Animal Care and Use Committee of the Medical University of South Carolina. Six-to-eight-week-old female Nu/J mice were housed with a temperature-controlled and 12-hour light/12-hour dark cycle environment. Tumors of MCF-7 parental, TamR, TamR.shCtrl or TamR.shLINC00152 cells were developed by injecting 1 × 10^7^ cells subcutaneously near mammary fat pads. Slow-release estradiol pellets (0.36 mg, 60 days; Innovative Research of America) were implanted subcutaneously (s.c) at the nape of the neck one day before injection. Tumor growth was regularly monitored, and size measurements were performed two times per week. Tumor volume was calculated using the formula (length × width^2^)/2. After MCF-7 TamR.shCtrl tumor size reached 100 mm^3^, the mice with shCtrl and shLINC00152 tumors were randomly divided into 2 groups, with 7-8 mice per group. Animals were treated with vehicle (corn oil) or tamoxifen (3 mg/kg in corn oil, by oral gavage) daily. The tumor size of each mouse was recorded twice a week. Mice were sacrificed 28 days after initiation of the treatment.

### Bioinformatics and statistical analysis

The microarray data sets, GSE6532^71^, GSE58644^72^, GSE42568^73^, GSE87411^74^ and GSE4922^75^ were download from the GEO database. The LINC00152 knockdown (KD) signature was generated using the cumulative list of genes down or upregulated upon LINC00152 knockdown using siLINC00152#1 and/or siLINC00152#2 in MCF-7 TamR cells. The KD score was then calculated by subtracting the sum of z-score of the genes upregulated from the sum of z-scores of the genes downregulated using the SPSS Statistics software. The ferroptosis gene signature comprises genes driving ferroptosis and was obtained from FerrDb V2 database^35^. The calcium signaling signature was obtained from KEGG database^41^, and the cAMP signature was generated using genes significantly upregulated (*P*-value < 0.001 and FC > 2) upon treatment of cancer cells with the cAMP analog, 8-CPT-cAMP for 24 hours^44^. All ferroptosis, calcium and cAMP scores were generated by summing up the z scores of the genes in each list, separately. The GSEA analyses in low LINC00152 KD score-expressing or low LINC00152-expressing patients were performed using the GSEA tool (Broad Institute).

The results are represented as mean□±□standard deviation (SD) or mean□±□standard error of the mean (SEM), as indicated in the figure legends. All statistical analyses were performed in GraphPad Prism Software. Comparisons between two groups were done using unpaired two-sided Student’s t-test. Comparison of the tumor volume graphs was done using two-way ANOVA with Dunnett’s multiple comparisons test. Survival curves were generated using Kaplan-Meier method, and significance between groups was calculated by Log-rank test. For correlation analysis, Pearson correlation coefficients were calculated. Experiments were repeated two to three times independently with similar results.

All other methods, including whole-transcriptome sequencing (RNA-seq) and data analysis, transient transfection with siRNAs, stable transfections using lentiviral vectors, quantitative RT-PCR analysis, mRNA stability assay using actinomycin, 3’UTR assay, Western blotting, in situ hybridization and immunohistochemistry are provided in Supplementary Methods. The Supplementary Tables for the sequences of shRNAs (**Supplementary Table S3**), siRNAs (**Supplementary Table S4**) and the list of antibodies (**Supplementary Table S5**) are also provided in the Supplementary Information file.

## Data availability

Gene expression data were downloaded from the NCBI Gene Expression Omnibus database under GSE1378, GSE6532, GSE58644, GSE42568, GSE87411 and GSE4922.

## Acknowledgements

We are thankful to the members of Ozgur Sahin laboratory for invaluable discussion and advice. We also thank the Flow Cytometry & Cell Sorting Shared Resource and Translational Science Laboratory of Medical University of South Carolina.

## Funding

This work was supported by research funding from American Cancer Society Research Scholar Grant RSG-19-194-01-CSM (O.S.), and in part, from the National Institutes of Health (NIH; R01CA267101 and R01CA251374 to O.S.), SmartState Endowment in Lipidomics and Drug Discovery (O.S.) and previously by TUBITAK Grant 215Z357 (O.S.), TUBITAK-CNRS Bilateral Grant (TUBITAK, 214S364 (O.S.) and CNRS (D.T.)), and Susan G. Komen Interdisciplinary Graduate Training to Eliminate Cancer Disparities (IGniTE-CD) GTDR17500160 (OzgeS). The core facilities utilized are supported by NIH (C06 RR015455), Hollings Cancer Center Support Grant (P30 CA138313) or Center of Biomedical Research Excellence (COBRE) in Lipidomics and Pathobiology (P30 GM103339).

## Contributions

Ozge S. designed and performed experiments, acquired and analyzed data, interpreted data, and prepared the paper; R.A. performed rescue experiments with PDE4D overexpression, lipid peroxidation and cAMP/Ca^2+^ measurement experiments and helped with shRNA generation; K-T. H. performed rescue experiments with LINC00152 overexpression; A.I. performed ISH of LINC00152 in ER+ breast cancer samples and contributed to data interpretation; M.U. performed the staining of PDE4D in ER+ breast cancer samples and contributed to data interpretation; N.B. contributed to cell viability and qRT-PCR experiments; P.G.E. contributed to bioinformatics analysis and determining LINC00152 expression in tumors; M.C. performed the RNA pulldown experiments; U.M.T. contributed to bioinformatic and experimental validations of LINC00152-PDE4D binding and analysis of the RNA-seq data; M.E.G. contributed to lipid peroxidation assay; H.B. was contributed to analysis of the differentially expressed lncRNAs in TamR cells; O. S. S. contributed to the in vivo experiment; Y.R. performed the RNA-seq and contributed to data interpretation; D.T. contributed to the bioinformatics analyses and data interpretation; D.G. contributed to the analyses of RNA-seq and data interpretation; B.O. contributed to the design of experiments and the interpretation of data; S.A, provided the clinical information of ER+ patients; A.U contributed to the staining of PDE4D and data interpretation and A.A. contributed to the ISH staining of LINC00152 in ER+ breast cancer samples and data interpretation; O.S. designed the study, oversaw experiments and data analyses, and prepared the paper. All authors reviewed and commented on the paper.

## Ethics statement

The study was approved by the Non□Interventional Clinical Research Ethics Committee of Hacettepe University (approval no: 2020/02-40). Informed consent was obtained from all patients.

## Competing interests

O.S. is the co-founder of OncoCube Therapeutics LLC, the founder and president of LoxiGen, Inc and the scientific advisory board member of A2A Pharmaceuticals. The other authors declare no potential conflict of interest.

