## Supplementary Information for "Targeting LINC00152 activates cAMP/Ca^2+^/ferroptosis axis and overcomes tamoxifen resistance in ER+ breast cancer"

Ozgur Sahin, PhD

Professor and SmartState Endowed Chair

Department of Biochemistry and Molecular Biology

Hollings Cancer Center

Medical University of South Carolina

86 Jonathan Lucas Street, Room HO712F,

Charleston, SC 29425

### SUPPLEMENTARY METHODS

#### Whole-transcriptome sequencing (RNA-seq) and data analysis

*RNA-sequencing of parental vs. TamR cells*

The RNA-seq of MCF-7 TamR vs parental cells was performed in triplicates using the Illumina HiSeq 2000 platform at McGill University and Genome Quebec Innovation Centre. Ribosomal RNA (rRNA)-depleted libraries were generated for each sample, and these sequences were multiplexed. RNA sequencing was performed for each condition (MCF-7 parental and MCF-7 TamR) in triplicates using the Illumina HiSeq 2000 platform at McGill University and Genome Quebec Innovation Centre. Around 65 million paired-end 2×100bp reads were generated for each replicate. FASTQC tool was used to check the quality of the sequencing data. Raw FASTQ sequences were aligned to the UCSC human

reference genome (hg19) using TopHat v2.1.0 with default parameters. *Cufflinks* was used to assemble and quantify transcripts from the mapped sequences. By utilizing reference genome annotation (UCSC hg19), *Cuffmerge* produced a single merged reference transcripts for differential expression analysis. For the identification of the differentially expressed genes, *Cuffdiff* was used with the default parameters. The 330 differentially expressed lncRNAs between parental vs. TamR cells (q value = 0.05) were retrieved and sorted based on the abundance (FPKM) in TamR cells (**Supplementary Data 1**).

*RNA-sequencing of TamR cells with LINC00152 knockdown*

The RNA-seq of MCF-7 TamR siControl, siLINC00152#1 and siLINC00152#2 was performed for each condition in duplicates at University of South Carolina Functional Genomics Core. FASTQC tool was used to check the quality of the sequencing data. Raw FASTQ sequences were aligned to the UCSC human reference genome (hg38) using TopHat v2.1.0 with default parameters. *Cufflinks* was used to assemble and quantify transcripts from the mapped sequences. By utilizing reference genome annotation (UCSC hg38), *Cuffmerge* produced a single merged reference transcripts for differential expression analysis. For the identification of the differentially expressed genes, *Cuffdiff* was used with the default parameters and the genes differentially expressed upon LINC00152 knockdown (siControl vs. siLINC00152#1 and/or siControl vs. siLINC00152#2) with the cut-off of 0.67 > FC > 1.5 and $P < 0.1$ were determined.

**Quantitative RT-PCR analysis**

Total RNA was extracted from cultured cells using TRIsure (Bioline), and cDNAs were generated using RevertAid RT Reverse Transcription Kit (Life Technologies). qRT-PCR analysis was performed with gene-specific primers using LightCycler 480 SYBR Green I Master kit (Roche). *HPRT1* and *ACTB* were used as housekeeping genes. The average Ct value was calculated from triplicates of each sample, and the relative mRNA expression was determined. Sequences of the qRT-PCR primers are listed in **Supplementary Table S2**.

**Stable transfections using lentiviral vectors**

An shRNA specifically targeting LINC00152 was designed using Vector Builder. The sequences are provided in **Supplementary Table S3**. To generate viral particles carrying shRNA vector, 6 μg of vectors along with the psPAX2 and pMD2.G packaging plasmids were co-transfected into HEK293FT cells in 6-well plate using lipofectamine (Invitrogen). After 48 hours of incubation, the viral particles were collected and transduced into MCF-7 TamR cells. Further selection was done by treating cells with the medium containing 2 μg/mL puromycin for 10 days. The PDE4D ORF expressing MCF-7 TamR.shLINC cells were generated by transfecting the MCF-7 TamR.shLINC cells with a PDE4D ORF vector (PDE4D cDNA (NM_006203.4) from GenScript), followed by selection with geneticin.

**Transient transfection with siRNAs**

siRNA transfections were done in P/S-free growth medium with reduced serum at a concentration of 40 nM using Lipofectamine 2000™ (Invitrogen) as previously described[1]. The list of siRNAs used (Dharmacon) are provided in **Supplementary Table S4**.

**mRNA stability assay using actinomycin**

To measure the changes in the stability of PDE4D mRNA upon LINC00152 knockdown, the TamR cells were transfected with 40 nM of siCtrl and siLINC00152. Two days later, cells were treated with 5 ug/mL actinomycin D and collected after 0, 60, 120, 240 minutes. RNA was extracted using TRIsure (Bioline), and cDNAs were generated using RevertAid RT Reverse Transcription Kit (Life Technologies). qRT-PCR analysis was performed using LightCycler 480 SYBR Green I Master kit (Roche). The fold change of PDE4D mRNA was measured by normalizing each time point to time zero for siCtrl and siLINC separately.

**3'UTR assay**

To test the binding of LINC00152 to *PDE4D* 3'UTR, cells were co-transfected with the LINC00152 overexpression vector together with *PDE4D* 3'UTR cloned in 3 different vectors. 48 hours after transfection, cells were lysed and luminescence signal was quantified using the Luciferase Assay System (Promega).

**Western blotting**

Western blotting was done as describe previously[2-4]. Briefly, total protein was extracted using RIPA buffer (150 mmol/L NaCl, 50 mmol/L Tris base pH 8.0, 1 mmol/L EDTA, 0.5% sodium deoxycholate, 1% NP40, 0.1% SDS, 1 mmol/L DTT, and 1 mmol/L Na3VO4) supplemented with Complete Protease Inhibitor (Roche). Protein concentrations were measured using BCA Protein Assay (Thermo Scientific). Equal amounts of protein lysates (15–20 μg) were separated on a 10% SDS-PAGE, transferred onto polyvinylidene difluoride membrane (Bio-Rad) and incubated with primary antibodies listed in **Supplementary Table S5**. The blots were developed using enhanced chemiluminescence (ECL) detection kit (Amersham Biosciences) after incubation with horseradish peroxidase–conjugated secondary antibody. β-Actin was used as a loading control.

**In situ hybridization and Immunohistochemistry**

LINC00152 RNA in situ hybridization was performed manually by RNAscope™ Probe- Hs-LINC00152 (Cat No. 464401, Advanced Cell Diagnostics (ACD), Hayward, CA) and RNAscope® 2.5 HD Detection Reagent – BROWN kit (Cat. No: 322310, ACD) in formalin-fixed paraffin-embedded (FFPE) tissues as defined in vendor's instructions. Briefly, 4-micron-thick unstained sections from FFPE blocks were prepared and deparaffinization with xylene and hydration with graded ethanol washes (96%, 90%, and 70%, respectively) was done. After endogenous peroxidase blockage, signal amplification and background suppression were performed as recommended in manual. Signals were developed with DAB and haematoxylin was used for counterstaining.

For the IHC staining of PDE4D protein in patient tumor samples from the Hacettepe cohort, antigen retrieval was done for 10 min with the EDTA buffer, and slides were incubated with the PDE4D antibody (Proteintech, clone 12918) at a dilution of 1:100 for 20 minutes at room temperature using Leica BOND-MAX/ISH automated immune-stainer.

The IHC staining of PDE4D in the FFPE slides from xenograft tumors was performed by deparaffinization at 60 $^oC$ for 1 hour, followed by rehydration in citrisolv for 5 min (3 times), 100% ethanol for 5 min (twice), 95% ethanol for 5 min (twice), deionized water for 5 min (twice). Antigen retrieval was done with Tris-EDTA pH=9 at 96 $^oC$ for 15 min, followed by cooling down to room temperature for 30 min, and washing with TBST for 5 min. Blocking was done at room temperature with Buffer W (IBA-Lifesciences) for 30 min. Then, the slides were incubated with the PDE4D antibody (Proteintech, clone 12918) at a dilution of 1:100 for 20 minutes at room temperature.

**SUPPLEMENTARY FIGURES**

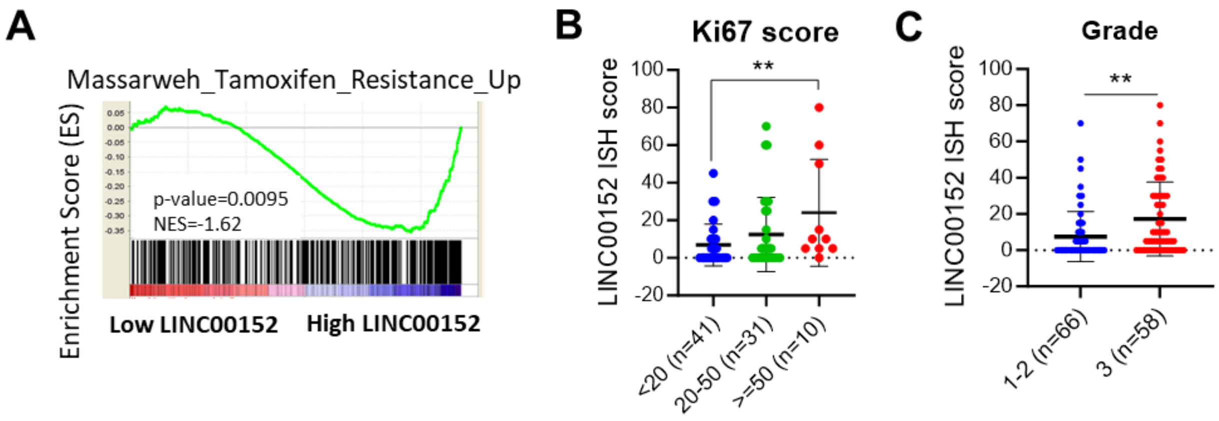


**Supplementary Figure 1. High LINC00152 expression correlates with tamoxifen resistance, higher proliferation and grade of ER+ patients treated with tamoxifen. A** GSEA analysis of genes upregulated in tamoxifen resistance showing enrichment in patients expressing high LINC00152. **B** The LINC00152 ISH score in ER+ breast cancer patients from the Hacettepe cohort with low (Ki67 score < 20), moderate (20 < Ki67 score < 50) and highly (Ki67 score > 50) proliferative tumors. **C** The LINC00152 ISH score in ER+ breast cancer patients from the Hacettepe cohort with low (1-2) vs. high (3) grade tumors. Data are presented as mean values ± standard deviation (SD). *P*-value for A was generated using the GSEA software (Broad Institute) and *P*-values for B and C were calculated with the unpaired, two-tailed Student's t test. ** *P*<0.01. NES: normalized enrichment score. Source data for this figure are provided as a Source Data file.

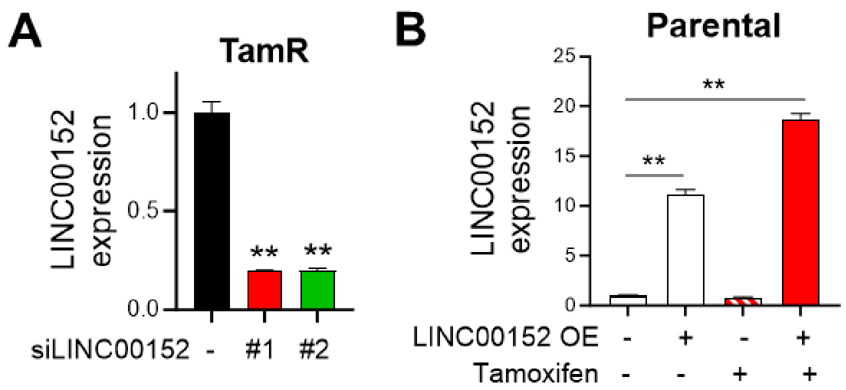


**Supplementary Figure 2. LINC00152 expression upon its knockdown or overexpression in ER+ breast cancer cell lines. A** qRT-PCR analysis of LINC00152 in siLINC00152-transfected MCF-7 TamR cells transfected with two different siRNAs. **B** qRT-PCR analysis of LINC00152 in T47D cells overexpressing LINC00152 and treated with or without tamoxifen. Data are presented as mean values ± standard deviation (SD). *P*-values were calculated with the unpaired two-tailed Student's t test. ** $P < 0.01$, n.s. not significant. Source data for this figure are provided as a Source Data file.

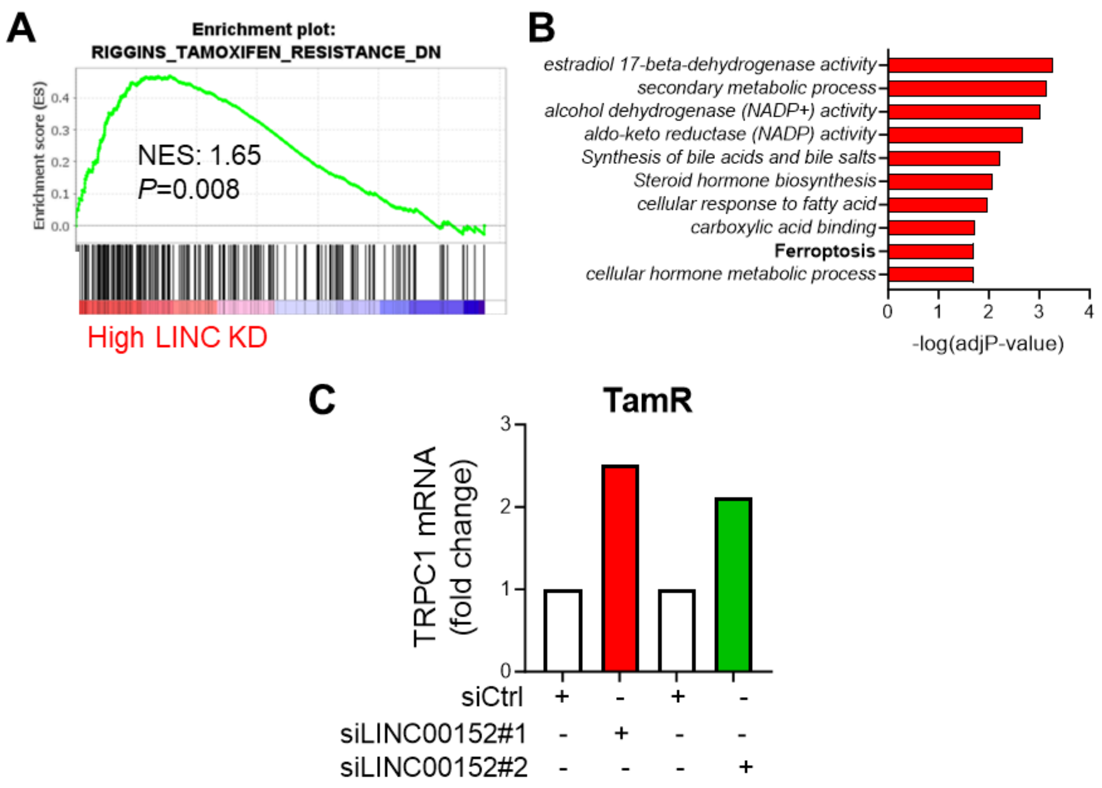


**Supplementary Figure 3. Association of LINC00152 knockdown (KD) score with tamoxifen resistance, ferroptosis, and TRPC1 upregulation upon LINC00152 knockdown. A** GSEA analysis in ER+ breast cancer patients showing enrichment of genes downregulated in tamoxifen resistance among patients expressing high LINC00152 knockdown (KD) score. NES: 1.65, *P* value = 0.008. **B** Pathway enrichment analysis among the differentially expressed genes in siLINC00152-transfected MCF-7 TamR cells. **C** TRPC1 fold change in MCF-7 TamR cells transfected with two different siRNAs against LINC00152 as determined by RNA-seq. *P*=0.023 and 0.042 for siCtrl vs. siLINC00152#1 and siCtrl vs. siLINC00152 #2 comparisons, respectively. Source data for this figure are provided as a Source Data file.

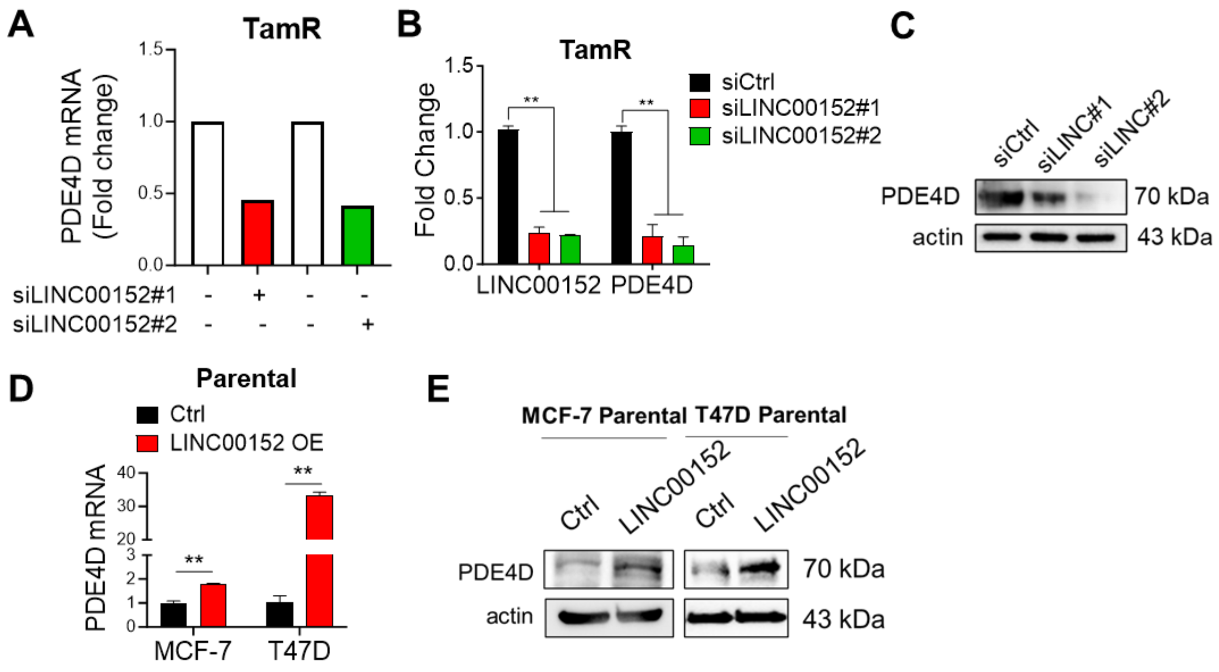


**Supplementary Figure 4. LINC00152 inhibition reduces PDE4D expression while its overexpression increases PDE4D, and PDE4D overexpression reverses cAMP signaling activity and ferroptosis under LINC00152 knockdown. A** PDE4D fold change in MCF-7 TamR cells transfected with two different siRNAs against LINC00152 as determined by RNA-seq. **B** qRT-PCR analysis of LINC00152 and PDE4D in MCF-7 TamR cells transfected with two different siRNAs against LINC00152. **C** Western blot analysis of PDE4D in MCF-7 TamR cells transfected with two different siRNAs against LINC00152. **D, E** qRT-PCR (D) and Western blot (E) analysis of PDE4D in parental MCF-7 and T47D cells overexpressing LINC00152. Data are presented as mean values ± standard deviation (SD). *P*-values were calculated with the unpaired, two-tailed Student's t test. ** $P < 0.01$. Source data for this figure are provided as a Source Data file.

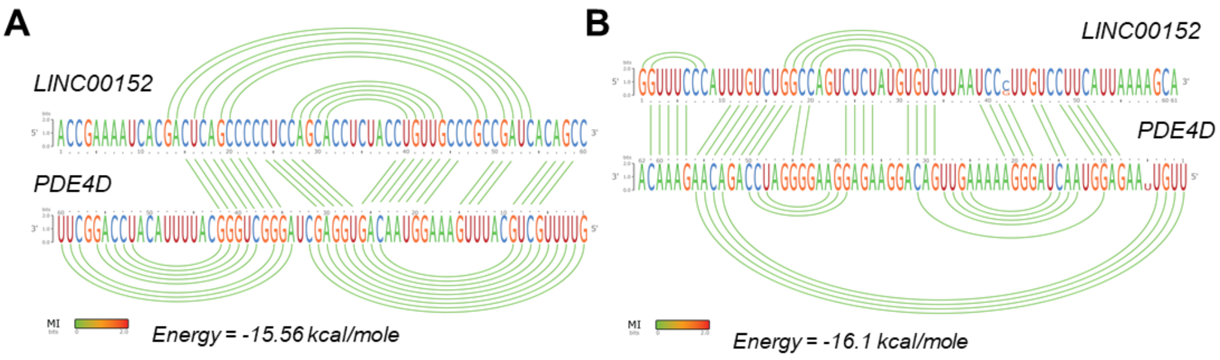


**Supplementary Figure 5. Predicted interactions between LINC00152 and *PDE4D* 3'UTR. A, B** Two of the predicted binding sites between LINC00152 and *PDE4D* 3'UTR as predicted by IntaRNA database. The visualization of the interaction was obtained using RILogo (https://rth.dk/resources/rilogo/). No mutual interaction (MI) between the bases has been detected. The predicted binding energies were provided. Source data for this figure are provided as a Source Data file.

### SUPPLEMENTARY TABLES

**Supplementary Data 1.** Significantly differentially expressed lncRNAs in MCF-7 TamR cells compared to parental counterpart.

Supplementary Data 1 is provided as a separate excel file.

**Supplementary Table S1.** The sequence of LINC00152 probe for RNA pull-down.

| ***Sequence of LINC00152 probe for RNA pulldown*** | | | |
|---|---|---|---|
| **Gene Name** | **NCBI Gene ID** | | **Probe sequence** |
| LINC00152 | 112597 | | 5'-CACCTTCCCAGGAACTGTGCTGTGAAGATCTGAAGACAGGC-3' |

**Supplementary Table S2.** The sequence of qRT-PCR primers.

| ***Sequences of qRT-PCR primers*** | | | |
|---|---|---|---|
| **Gene Name** | **NCBI Gene ID** | | **Primer sequence** |
| ACTB | 60 | Forward | 5'-CCAACCGCGAGAAGATGA-3' |
| | | Reverse | 5'-CCAGAGGCGTACAGGGATAG-3' |
| HPRT | 3251 | Forward | 5'-TGACCTTGATTTATTTTGCATACC-3' |
| | | Reverse | 5'-CGAGCAAGACGTTCAGTCCT-3' |
| PDE4D | 1612 | Forward | 5'-ATTGCCAGTGATATACACGGA-3' |
| | | Reverse | 5'-GTTTCGTACAGTTCGCAGAC-3' |
| LINC00152 | 112597 | Forward | 5'-GAGCCACCAGCCTCTCCTTG-3' |
| | | Reverse | 5'-GGGCTGAGTCGTGATTTTCG-3' |

**Supplementary Table S3.** The sequence of shRNA against LINC00152.

| **Gene Name** | **NCBI Gene ID** | **Company** | **Vector Name** | **Sequence** |
|---|---|---|---|---|
| LINC00152 | 112597 | Vector Builder | pLV[shRNA]-EGFP:T2A:Puro-U6>(sh-CYTOR-4) | CAGGAAGCTCTATGACACATTCAAGAGATGTGT CATAGAGCTTCCTG |

**Supplementary Table S4.** Sequences of siRNAs.

| ***Sequences of siRNAs used*** | | | | |
|---|---|---|---|---|
| **Gene Name** | **NCBI Gene ID** | **Company** | **Catalog number** | **Sequence** |
| LINC00152#1 | 112597 | Dharmacon | N-015149-20-005 | GUCCUUCAUUAAAAGCAAA |
| LINC00152#2 | 112597 | Dharmacon | N-015149-18-005 | ACAGGAAGCUCUAUGACAC |

**Supplementary Table S5.** List of antibodies used in Western blot (WB) experiments.

| **Antibody** | **Provider** | **Catalog number** | **WB dilution** |
|---|---|---|---|
| Beta-actin | MP Biomedicals | 691001 | 1:10000 |
| PDE4D | ProteinTech | 12918-1-AP | 1:1000 |
| GPX4 | Cell Signaling | 52455 | 1:1000 |
| xCT | Cell Signaling | 12691 | 1:1000 |
| FTH1 | Cell Signaling | 4393 | 1:1000 |
| p-PKA (Thr197) | Cell Signaling | 4781 | 1:1000 |
| p-CREB (S133) | Cell Signaling | 9198 | 1:1000 |
| PKA | Cell Signaling | 5842 | 1:1000 |
| CREB | Cell Signaling | 4820 | 1:1000 |
| TRPC1 | Santa Cruz | sc-133076 | 1:1000 |
